# Sleep spindles track cortical learning patterns for memory consolidation

**DOI:** 10.1101/2021.09.01.458569

**Authors:** Marit Petzka, Alex Chatburn, Ian Charest, George M. Balanos, Bernhard P. Staresina

## Abstract

Memory consolidation, the transformation of labile memory traces into stable long-term representations, is facilitated by post-learning sleep. Computational and biophysical models suggest that sleep spindles may play a key mechanistic role for consolidation, igniting structural changes at cortical sites involved in prior learning. Here we tested the resulting prediction that spindles are most pronounced over learning-related cortical areas and that the extent of this learning-spindle overlap predicts behavioural measures of memory consolidation. Using high-density scalp Electroencephalography (EEG) and Polysomnography (PSG) in healthy volunteers, we first identified cortical areas engaged during a temporospatial associative memory task (power decreases in the alpha/beta frequency range, 6-20 Hz). Critically, we found that participant-specific topographies (i.e., spatial distributions) of post-learning sleep spindle amplitude correlated with participant-specific learning topographies. Importantly, the extent to which spindles tracked learning patterns further predicted memory consolidation across participants. Our results provide empirical evidence for a role of post-learning sleep spindles in tracking learning networks, thereby facilitating memory consolidation.

## Introduction

Sleep after learning bolsters memory retention, a process referred to as sleep-dependent memory consolidation ^1–3^. In recent years, sleep spindles – transient 12-15 Hz oscillations generated within thalamo-cortical loops - have emerged as a prime mechanistic vehicle to support consolidation ^4–9^. Previous studies have linked spindles to consolidation in terms of their density (the number of discrete spindle events per minute) ^10^, power ^11^ and activity, a combination of duration and amplitude ^12^. Despite their ubiquity, however, the specific role spindles play for memory consolidation remains poorly understood.

Ultimately, effective learning requires structural brain changes, beginning at the synaptic level ^13,14^. A hallmark computational/biophysical framework ^15^ suggests that spindles are particularly well-suited to induce changes in synaptic plasticity. Specifically, spindles gate influx of calcium (Ca^2+^) into pyramidal dendrites, setting early synaptic consolidation processes in motion. Empirical support for this model has been provided by in-vitro application of spindle-like firing patterns ^16^ as well as by showing a direct modulation of Ca^2+^ activity in cortical pyramidal dendrites as a function of spindle power during natural sleep in rodents ^17^. Moreover, cortical microelectrode array recordings in humans have shown that spindles group co-firing of single units within 25 ms, i.e. within a time window conducive to spike-timing-dependent plasticity ^18^. Critically, however, in order for spindles to promote memory consolidation in an adaptive fashion, they need to show some degree of regional specificity. That is, not only would global synaptic consolidation be of limited use in an ever-changing landscape of tasks, but it would also be at direct odds with extant models emphasising the role of sleep in global synaptic downscaling ^19^. Instead, adaptive consolidation has to be selective, specifically strengthening local circuits involved in prior learning.

The current study thus set out to assess (i) whether spindles track specific pre-sleep learning patterns and (ii) whether this learning-spindle overlap supports memory consolidation. To this end, we employed a demanding memory task giving rise to rich and idiosyncratic activation patterns during encoding (*Memory Arena*, Figure 1). After learning, participants took a 2-hour nap before their memory retention was tested. This protocol allowed us to examine whether spindles recorded during this nap would track participant-specific learning patterns and whether the extent of this learning-sleep overlap would predict behavioural expressions of consolidation.

**Figure 1.**
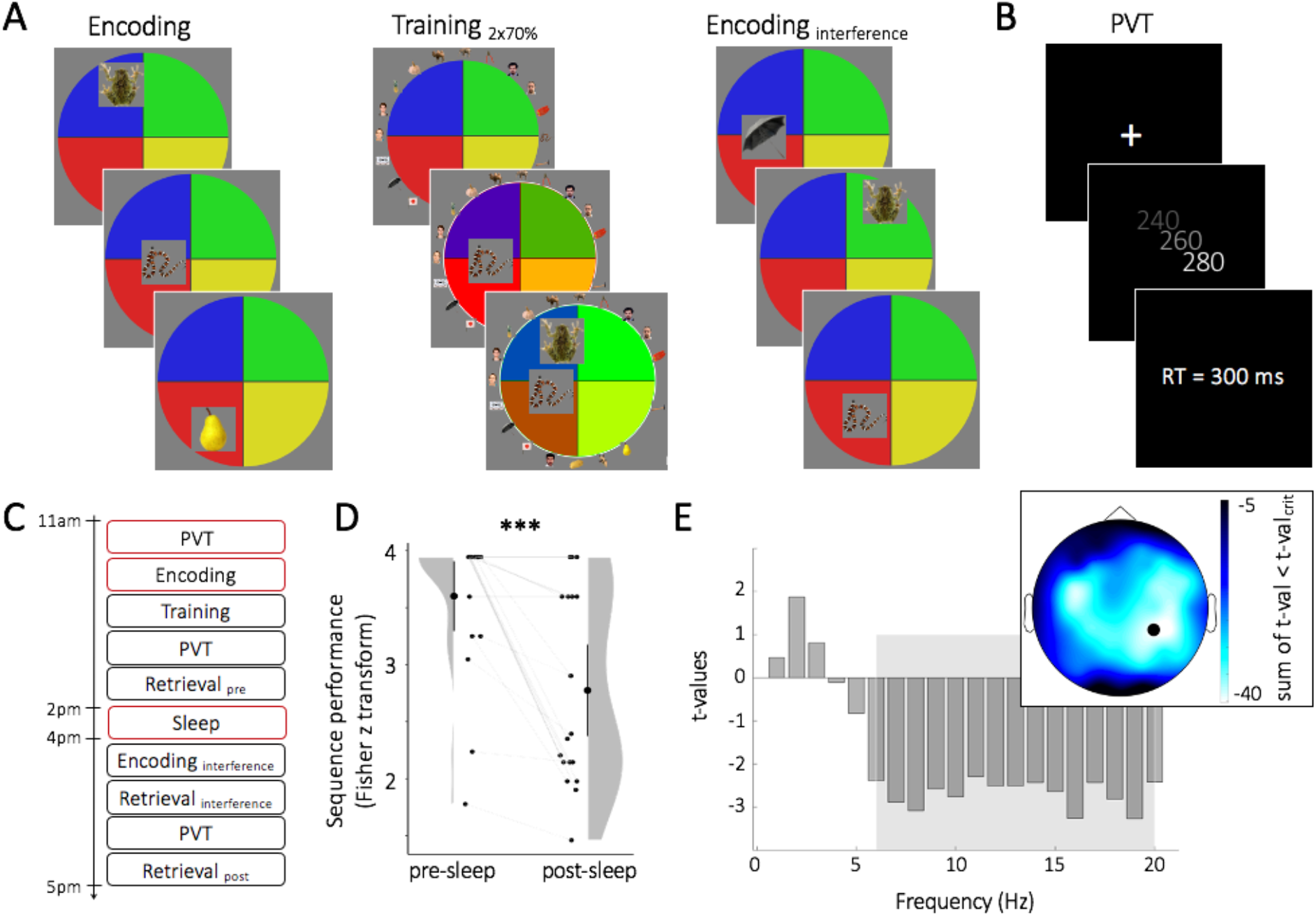
Task, experimental design, behavioural results and group EEG encoding pattern. (A) *Memory Arena.* During encoding 20 objects were presented in a specific sequence at different spatial positions. Both sequence and spatial position had to be encoded. Training (and retrieval) started with all 20 objects arranged around the arena and participants had to drag and drop the objects in the correct sequence to the correct spatial position. During training, feedback was given after each trial and errors were corrected. Training was completed after reaching a performance criterion of 70% twice in a row. Retroactive interference was induced by encoding the same objects but in a different sequence and at different spatial positions. (B) Every trial of the Psychomotor Vigilance Task (PVT) started with a fixation cross. After a delay of 8-15 sec a counter started. Participants had to press a key as fast as possible and received feedback about their reaction time (RT). (C) Participants performed the PVT, encoding, training and first retrieval before the 2-hour nap. After the nap, an interference session was employed (1x encoding and retrieval, no training), followed by the second retrieval of the originally learned arena. (D) Sequence memory significantly decreased from pre-to post-sleep. Single participant data, density plots and group means with 95% CIs are shown. *** = p < .001. (E) Comparison of oscillatory power during Memory Arena vs. PVT (thresholded at p < .05 cluster corrected), revealing a significant power decrease from 6 – 20 Hz during encoding (grey rectangle), most pronounced over temporo-parietal areas (bars shown for electrode CP4 - black circle on topography plot).

## Results

### Behavioural results

We employed a recently developed memory paradigm called ‘Memory Arena’ 20 in which participants learn the temporospatial arrangement of objects in a circular enclosure across multiple training rounds (see Figure 1A and Methods for details). Memory for the temporospatial arrangement was assessed in a first retrieval block (pre-sleep retrieval) which was followed by a 2-hour nap (see Table S1 for descriptive data of sleep stages). Following the nap, participants were instructed to learn a new temporospatial arrangement of the same objects (retroactive interference), after which they were asked to retrieve the original arrangement (post-sleep retrieval, Figure 1C). As the dependent measure, we use sequence memory performance as our previous work suggested this was the measure most sensitive to capture sleep-dependent consolidation. As expected, we found a significant decrease in sequence performance from pre-to post-sleep retrieval of the original sequence (t(18) = 4.65, p < .001, Figure 1D). For further analyses, memory consolidation is defined as memory retention, i.e., the relative change in sequence performance from pre-to post-sleep retrieval.

### EEG results: Spindle amplitude tracks encoding patterns

To assess whether sleep spindles track learning sites, we first derived an ‘encoding pattern’ for each participant. To specifically unravel learning-related activity, we contrasted oscillatory power (1-20 Hz) during encoding with power during a control condition (Psychomotor Vigilance Task, PVT). On the group level, this contrast revealed a significant power decrease in the alpha/beta frequency range (6-20 Hz) during encoding relative to the PVT, particularly over right temporo-parietal areas (Figure 1E). This result is consistent with previous findings linking decreases in alpha/beta power to memory processes ^21–25^. The reliable group effect notwithstanding, there was considerable variability in participant-specific effect topographies of the 6-20 Hz power decrease (Figure S1), allowing us to explore whether these participant-specific encoding patterns would bias particular event characteristics during subsequent sleep.

As outlined in the introduction, we hypothesized that the topography (i.e., spatial distribution) of sleep spindles might be modulated by engagement during pre-sleep learning. We thus algorithmically detected sleep spindles during the post-learning nap (see Methods) for every channel and extracted their amplitude as well as duration and density. As a control, we performed the same analyses for algorithmically detected slow oscillations (SOs), which have also been linked to memory consolidation ^11,26,27^. At the group level, spindles and SOs showed the established prevalence over centro-parietal and frontal areas, respectively (see Figure S2 for amplitude, density and duration topographies of spindles and SOs). Furthermore and in line with previous observations ^28–31^, detected spindle events were on average temporally coupled to the up-state of the SO signal (0.3-1.25 Hz, see Figure S3).

We next turned to the question whether inter-individual differences in the topography of sleep events relate to inter-individual differences in learning topographies. For each participant, the encoding topography (6-20 Hz power relative to the PVT across 58 channels) was correlated (Spearman’s rho) with the corresponding topography of 6 different sleep patterns (amplitude, duration and density for spindles and SOs across 58 channels). Note that all sleep measures are positively scaled except the SO amplitude. For simplicity, we unified all scales by taking the absolute value of the SO amplitude. As encoding activity is associated with a decrease in power (negatively scaled values), encoding-sleep overlap would be signified by negative correlations.

Participant-specific correlation values were then evaluated at the group level. First, we conducted a 2 (event type: spindles vs. SOs) x 3 (event characteristic: amplitude, duration, density) repeated measures ANOVA (Figure 2A), assessing whether particular sleep event topographies track idiosyncratic encoding topographies. Results revealed that topographies of overall spindle characteristics correlated with encoding patterns to a greater extent than SO characteristics (main effect of event type F(1,18) = 13.12, p <.001, Figure 2B). This was the case for event amplitudes (t(18) = −3.16, p = .005), durations (t(18) = −2.53, p = .021) as well as densities (t(18) = −2.14, p = .046). Furthermore, the correlation between all three spindle characteristics with the encoding pattern was significantly smaller than 0 (spindle amplitude: meanr = −0.38, t(18) = −3.50, p = .003, spindle duration: meanr = −0.27, t(18) = −3.26, p = .004, spindle density: meanr = −0.30, t(18) = −3.56, p = .002). There was a trend for the main effect of event characteristic (amplitude topographies correlating strongest with encoding patterns, F(2,36) = 2.61, p = .078) and no interaction (F(2,36) = 0.63, p = .532).

**Figure 2.**
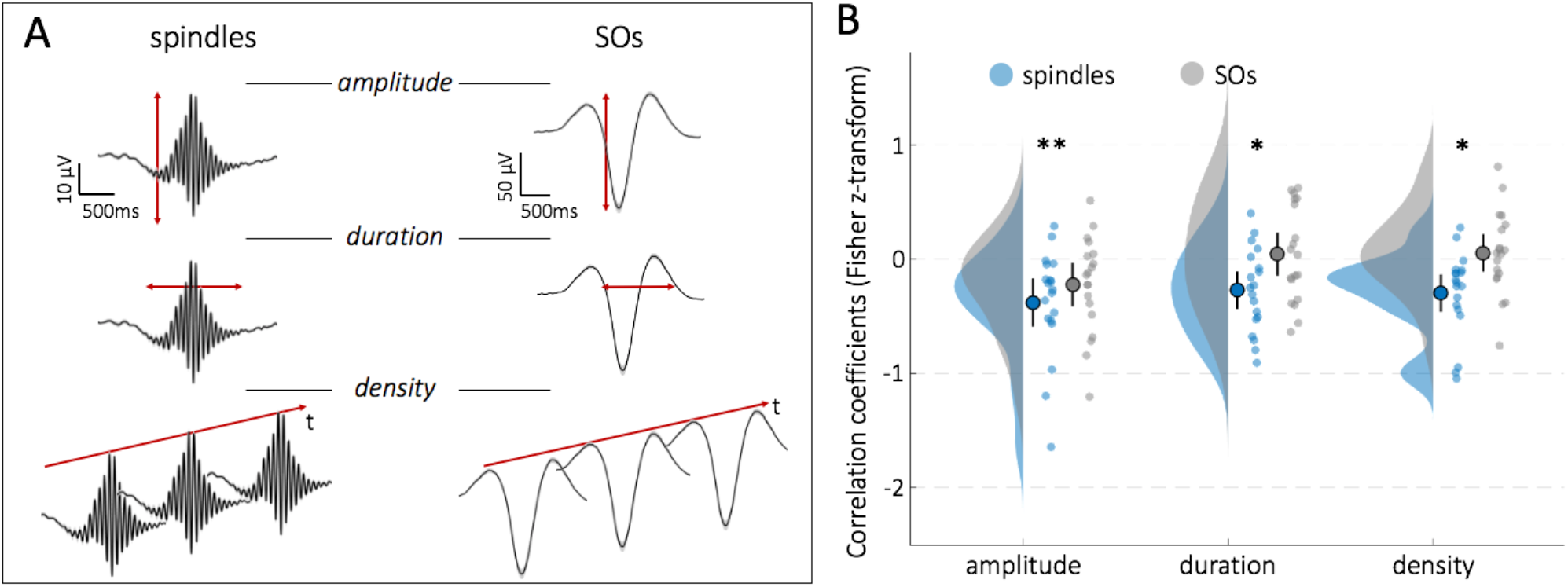
Sleep spindles track cortical learning sites. (A) Grand average (mean ± 95% CI) of spindles and SOs at electrode position Cz. Amplitude, duration and density were extracted for each participant and channel. (B) Across participants, encoding topographies were significantly more strongly correlated with topographies of spindles than SOs (density plots, group means, 95% CIs and single participant data are shown. ** = p < .01. * = p < .05).

To account for the possibility that spindle amplitude, duration and density topographies are correlated amongst each other, we conducted follow-up partial correlations between encoding patterns and spindle characteristics. Interestingly, the link between encoding and spindle amplitude remained significantly different from 0 when partialling out spindle duration or spindle density (for both: meanr < −0.23, t(18) < −2.11, p < .049). When partialling out spindle amplitude, however, the overlap between spindle duration/density and encoding pattern was not significantly different from 0 (duration: meanr = −0.15, t(18) = −1.82, p= .086; density: meanr = −0.01, t(18) = −0.15, p = .885). Together, these results suggest that the spindle-encoding overlap is predominantly driven by spindle amplitude.

To examine whether the overlap with encoding patterns might be restricted to spindles that are coupled to SO up-states, we directly compared the encoding-spindle overlap (amplitude topographies) for spindle events with higher vs. lower coupling (see Methods). We observed no significant difference between the two event types (t(18) = −0.66, p = .521, see Figure S3C).

Finally, we tested whether the overlap of sleep spindle amplitude and encoding activation is linked to behavioural expressions of memory consolidation. To this end, we correlated the encoding-spindle amplitude overlap with the relative change in sequence performance from pre-to post-sleep retrieval (memory retention) across participants. Indeed, a significant negative correlation was observed (r = −0.58, p = .010), indicating that participants who showed greater retention of sequence memory also had a greater overlap (signified by a more negative value) between encoding and sleep spindle topography (Figure 3A). To ensure that the link with behaviour was driven by participant-specific encoding-spindle overlap, we first shuffled the encoding topographies between participants while retaining participant-specific sleep spindle topography and behavioural performance. Likewise, we shuffled the sleep spindle topographies between participants while retaining participant-specific encoding topography and behavioural performance (see Figure 3B for visualization). That way we generated a distribution under the null hypothesis that (a) the encoding topography or (b) sleep spindle topography is irrelevant for the observed correlation with behaviour. As shown in Figure 3C, the empirical correlation between encoding-spindle overlap and behaviour significantly exceeded the null distributions in both cases (p = .011 for (a) and p = .018 for (b) based on 1000 permutations).

**Figure 3.**
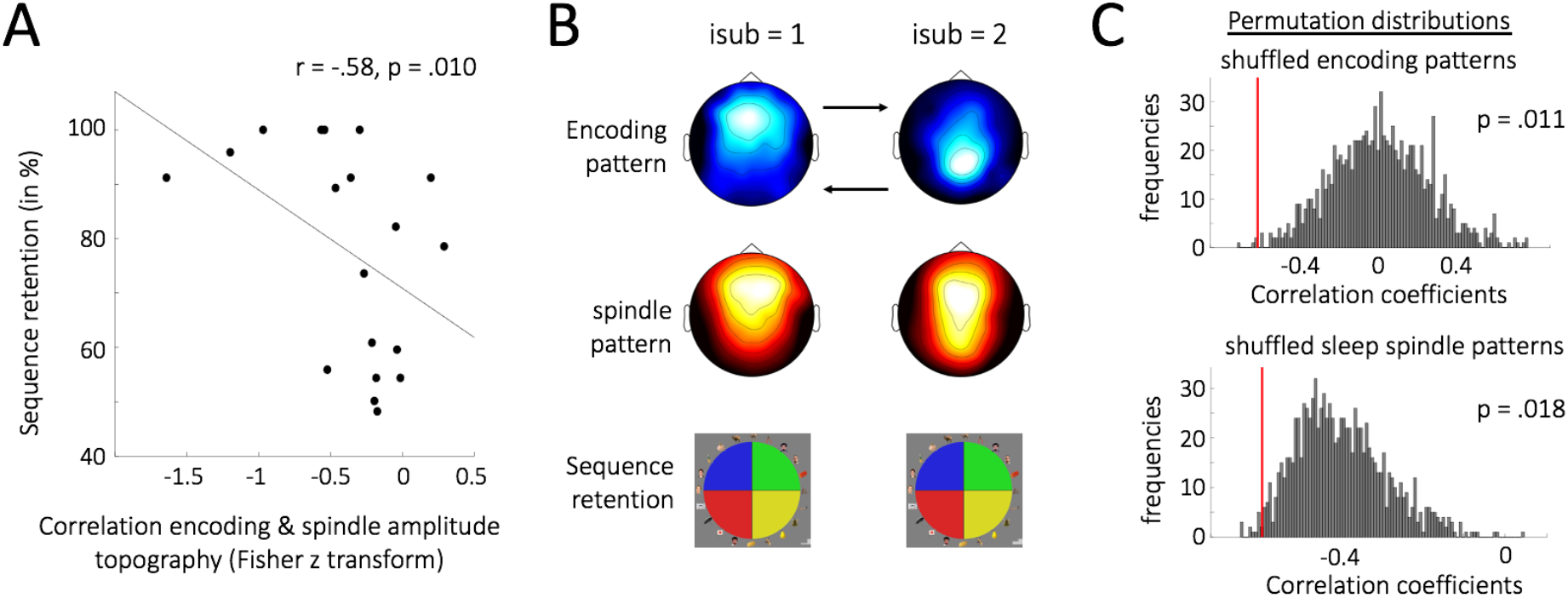
Extent of encoding-spindle overlap predicts memory consolidation. (A) The correlation between encoding and spindle amplitude topographies is predictive for sequence retention. The more negative the correlation between spindle amplitude and encoding topographies (i.e., the greater the overlap), the greater the levels of sequence retention across sleep. (B) Schematic for generating the null distributions in C. Encoding or sleep spindle topographies were shuffled across participants while the respective other topography (spindle amplitude or encoding power) and behavioural performance were retained. (C) The observed correlation between encoding-spindle amplitude overlap and sequence retention exceeds both null distributions of encoding or spindle pattern exchangeability.

## Discussion

Despite accumulating evidence linking sleep spindles to memory consolidation ^5–8,10–12,32^, their specific function has remained elusive. Here we tested predictions derived from biophysical/computational models, i.e., that spindles are preferentially expressed over learning-related sites where they might induce early stages of synaptic plasticity ^15^. Using a recently developed paradigm sensitive to sleep-dependent memory consolidation ^20^ and geared towards eliciting rich idiosyncratic encoding patterns (Figure 1), we first demonstrate that sleep spindles – in particular the topography of spindle amplitudes – track cortical patterns of memory encoding (Figure 2). This overlap of encoding patterns with spindles was significantly stronger than that with slow oscillations (SOs), ruling out spurious correlations driven by generic signal properties across EEG channels. Importantly, we additionally reveal a functional link between the observed encoding-spindle overlap and memory consolidation, expressed in greater overlap associated with greater levels of memory retention (Figure 3).

Previous work has revealed a number of spindle characteristics that render them well-suited for inducing local plasticity in task-dependent/learning-related brain regions. For instance, intracranial recordings in humans have shown that sleep spindles are local rather than global ^33,34^ and scalp EEG/MEG has demonstrated high levels of inter-subject variability of spindle topographies ^32,35^. Moreover, spindle topographies vary as a function of prior learning tasks at the group level – spindles over left frontal areas are related to consolidation of verbal material ^36^, whereas spindles over parietal areas are related to consolidation of visuospatial memories ^37^. However, as brain activity was not measured during learning in these studies, the link between learning activation and post-learning spindle topography remained conjectural. Another study employed simultaneous EEG-fMRI recordings and found a BOLD increase in learning-related ventro-temporal regions time-locked to sleep spindles at electrode position Cz ^38^, but whether spindles per se would be preferentially expressed at learning-related sites remained unclear due to limited EEG coverage. Finally, we recently showed that learning content can be decoded in the presence of centrally recorded sleep spindles ^30,39^, but decoding was based on raw EEG data rather than on spindle topographies. In short, despite converging evidence that spindle expression can be local, idiosyncratic and flexible, we here provide first evidence that they track participant-specific learning patterns in service of memory consolidation. These results dovetail with a recent computational study showing that spindles promote independent reactivation of multiple memories at network locations corresponding to awake training ^40^.

It deserves mention that the spatial resolution afforded by scalp EEG is relatively coarse, only capturing macro-scale topographies of brain networks. Intracranial recordings/Electrocorticography (ECoG) would provide much finer resolution, albeit at the expense of comprehensive whole-brain coverage and consistency across participants. That said, a recent study used cortical microelectrode array recordings (Utah Arrays) in four pre-surgical epilepsy patients and demonstrated different spindle- and unit firing dynamics across a 10 x 10 electrode grid covering < 15 mm^2 18^. This suggests that spindle deployment might be sufficiently fine-tuned in space to selectively strengthen local microcircuits of learning networks.

A key open question is how exactly the deployment of spindles to learning-related cortical sites is governed. One speculative possibility is that circuit-specific encoding activation establishes transient synaptic tags. Spindles are initially broadcast widely and stochastically during sleep but resonate more strongly when coinciding with those synaptically primed circuits. This leads to higher spindle amplitudes and a concomitant increase in Ca^2+^ influx, completing the tag-and-capture cycle suggested to underlie long-term-potentiation ^41^. Increased spindle amplitude, as observed here and in other studies ^38,42,43^, would thus reflect elevated local neural co-activation as a vestige of prior task engagement.

Another possibility is that the same thalamic circuits control deployment of attentional resources during wake task performance and spindles during sleep. For instance, a recent study capitalised on the orientation-specific response potentiation (OSRP) in mouse primary visual cortex (V1), which reflects enhanced firing to a visual grating of particular orientation several hours after initial exposure/training. Importantly, neurons in the lateral geniculate nucleus (LGN) of the thalamus already showed orientation-selective tuning during and immediately after training. During post-training sleep, thalamocortical coherence mediated by sleep spindles drove post-sleep orientation-selective tuning in V1 ^44^. Additional work is needed to elucidate whether similar mechanisms apply to more complex tasks in humans, but accumulating evidence across species has linked thalamic microcircuits to a wide range of cognitive tasks ^45^. This scenario, in which thalamocortical dynamics during learning bias the path of spindle deployment during sleep, is reminiscent of models of hippocampal functioning. Specifically, according to the hippocampal indexing theory ^46^, the hippocampus retains pointers to cortical circuits involved in learning. Upon presentation of a partial cue, hippocampus drives reinstatement (pattern completion) in cortical target sites. Recent work in rodents ^47^ and humans ^48^ points to a cortical-hippocampal-cortical loop around hippocampal ripples, and a tentative scenario might be that the initial cortical response in this loop is mediated by the aforementioned thalamo-cortical spindle projections.

Apart from spindles, memory processing during sleep has been linked to slow oscillations (SOs) and delta (1-4 Hz) rhythms, together referred to as slow wave activity (SWA) ^6,19,49,50^. One seminal study showed that SWA was specifically increased over central cortical areas thought to be involved in prior motor learning ^27^. Likewise, wake immobilisation of a participant’s arm led to a decrease in SWA over corresponding motor areas ^51^. Interestingly though, analogous effects were seen in the spindle/sigma band in both studies, raising the possibility that both SOs/SWA and spindles contribute to sleep-dependent consolidation. One pressing question is whether consolidation relies on concomitant or on sequential occurrence of these two sleep events. Speaking to the importance of concomitant SWA-spindles, a recent rodent study showed that Ca^2+^ activity was increased threefold when spindles were coupled to slow oscillations ^52^. The importance of coupled SO-spindle complexes has been further corroborated by a series of recent findings linking the precision of SO-spindle coupling to memory function in ageing ^28,29^ and to reinstatement of prior learning experiences during sleep ^30^. However, we did not observe greater overlap of encoding patterns with spindles coupled vs. not coupled to SOs in the current study (Figure S3C). Interestingly, a recent EEG/MEG study showed that the topography of spindles was unaffected by the topography of concurrent SOs ^32^. This raises the possibility that consolidation of learning patterns relies, at least in part, on the sequential occurrence of spindles and SOs. Indeed, in the original framework ^15^ as well as in a recent computational model ^40^, it is proposed that SOs, which show enhanced prevalence during later sleep stages, further potentiate strong synapses, incidentally leading to downscaling of weak synapses ^53^. In other words, sleep-dependent consolidation might rely on a multi-stage tagging and capture sequence, initiated by wake task performance, potentiated by thalamocortical sleep spindles in conjunction with hippocampal ripples, and completed by SOs. Whole-night recordings would be better-suited to test this notion than the current nap design.

To conclude, the present study demonstrates that sleep spindles track cortical areas engaged during prior learning and that the extent of learning-spindle overlap predicts levels of memory consolidation. An exciting avenue for future work will be to elucidate the spatiotemporal dynamics between spindles, ripples and slow oscillations across the hippocampus and neocortical sites, both in close temporal proximity and across a whole night of sleep.

## Acknowledgements

This work was supported by a Wellcome Trust/Royal Society Sir Henry Dale Fellowship (107672/Z/15/Z) to B.P.S.

## Author Contributions

Conceptualisation: M.P., A.C., B.P.S.; Methodology: M.P., I.C., B.P.S.; Investigation: A.C.; Formal Analysis: M.P., B.P.S.; Writing - Original Draft: M.P., B.P.S.; Writing – Review & Editing: M.P., A.C., B.P.S.; Visualisation: M.P., B.P.S.; Supervision: G.M.B., B.P.S.; Funding Acquisition: B.P.S.

## Methods

### Participants

22 participants were tested. Due to technical issues during data collection, 3 participants had to be excluded resulting in 19 participants for the final sample (mean_age_ = 20.7, range_age_ = 18-31, female = 15).

Pre-screening ensured that participants had no history of neurological or psychiatric disorders and a normal sleep-wake cycle. Participants were instructed to get up one hour earlier than normal and avoid caffeine the day of the experiment. After participating in the study, participants received monetary reimbursement. The study was approved by the University of Birmingham Research Ethics and Governance Committee and written informed consent was obtained from all participants before the start of the study.

### Paradigm and procedure

Memory Arena and PVT were implemented via custom scripts in MATLAB 2016a (MathWorks, Natick, USA). For the PVT, functions of the Psychophysics Toolbox Version 3.0.14 ^54^ were used.

#### Memory Arena

The *Memory Arena* consists of a circle divided into coloured quarters (upper left: blue, upper right: green, lower right: yellow, lower left: red). Within the circle, objects are sequentially presented in different spatial positions. Participants have to learn both the sequence in which the objects were presented as well as the spatial position of each object.

20 target objects (images of 5 faces, 5 natural objects, 5 animals and 5 manmade objects) were randomly selected from a stimulus pool of 40 objects (coloured and presented on a grey 90×90 pixels square) ^57,58^. The spatial position of each object was restricted by the position of other objects. Consequently, there was no overlap between objects, but objects possibly covered more than one colour wedge.

#### Psychomotor Vigilance Task (PVT)

A white fixation cross was presented in the middle of the screen. After on average 6 seconds (jitter ± 4s), a counter replaced the fixation cross. The counter started at 0 and counted forward in 20 ms steps to 2000. Upon start of the counter, participants had to press the space bar as fast as possible. After the key press, feedback about their reaction time was displayed for 2s (Figure 1B). Overall, the PVT lasted 2 minutes.

#### Procedure

The experimental session started at 10 am with the application of electroencephalography (EEG), electromyography (EMG) and electrooculography (EOG).

Approximately one hour later at 11 am, participants started with a short practice session (~20 sec) of the PVT which was followed by the actual task. Before they continued with the Memory Arena, participants received written instructions and performed a practice session (with 3 objects) of each Memory Arena part (encoding, training and retrieval). Participants were instructed to associate and combine the objects into a coherent story.

During the encoding part of the Memory Arena, all 20 objects were sequentially presented within the circle. Participants confirm processing of each object by clicking on it. The current object then disappeared, and the next object was presented.

Directly after the encoding part, a training session was conducted. The training session started with all 20 objects arranged around the arena. The objects had to be dragged and dropped in the correct sequence to their correct spatial position. If an error was made regarding the sequence or spatial position, the arena turned red and the error was corrected. Sequence errors were defined as object *i* not being placed at the *i^th^* position. Spatial errors were defined as the overlap between correct and chosen position of an object being less than 25%. After potential error corrections, the object remained at its correct spatial position and the next object had to be selected and placed in the arena. When all 20 objects were placed, feedback about the overall performance was presented. The overall performance was defined as the number of correct objects divided by the total number of objects, where an object was classified as correct when sequence as well as spatial position were correct. Participants finished training after reaching 70% overall performance in two consecutive runs.

A second PVT then followed the training session. After the PVT, the pre-sleep retrieval was completed. Like the training, the retrieval started with all 20 objects arranged around the arena which had to be dragged and dropped in the correct sequence to their correct spatial position. Importantly though, errors were not corrected and no feedback was provided.

Participants started the 2-hour nap between 1pm - 2.30pm (see Table S1 for descriptive sleep data). Following the nap, participants continued with an interference task. We were particularly interested in using an experimental design that is suitable to capture sleep-dependent memory consolidation. In a previous study, we used the same task (Memory Arena) and found sleep-dependent consolidation effects with a combination of a 2×70% training threshold (70% overall performance in two consecutive runs) and the induction of retroactive interference directly after sleep ^20^. Therefore, in this study, we applied the same methods.

During the interference task, participants had to encode the same 20 objects but in a different sequence and at different spatial positions (Figure 1A). The difference between the old and the new, interfering, spatial position of every object was at least 5 pixels (Euclidean distance). Encoding of the interfering positions was conducted in the same way as the original encoding. Subsequently, participants had to retrieve the interfering sequence and spatial positions (without prior training). Lastly, another PVT and the post-sleep retrieval (of the original sequence and spatial positions) were performed. Until the start of the post-sleep retrieval, participants were unaware of the final test.

### EEG data recording

EEG data were recorded using a Brain Products 64-channel EEG system and were sampled at a rate of 1000 Hz. Electrodes were arranged according to the 10-20 system (including FCz as reference, AFz as ground and left and right mastoids). Two electrodes were placed on the chin to record muscle activity (electromyography, EMG) and two electrodes recorded eye movements (electrooculography, EOG).

### Behavioural analysis

In a previous study using the same paradigm, we found that sequence performance was most sensitive measure to capture sleep-dependent memory consolidation ^20^. Consequently, all following analyses focus on sequence performance.

To calculate sequence performance, the selected order of all 20 object was correlated with a vector ranging, in ascending order, from 1 to 20. This correlation approach is preferable to simply counting the correct sequence position of every object, as it reflects both the correct sequence position of every object as well as correct transitions between objects. Correlation values were Fisher z-transformed for further statistical analyses. To test for a significant sequence performance change from pre-to post-sleep retrieval, a paired t-test was computed.

For correlating the change in sequence performance with the EEG data, a sequence retention score was calculated as the relative change from pre- to post- sleep sequence retrieval: 100*(post/pre).

### EEG analysis

EEG analyses were performed using the FieldTrip toolbox 59 and custom written scripts in MATLAB.

#### Encoding pattern

To remove eye movements from the data, an independent component analysis (ICA) was used. Data were down-sampled to 200 Hz, re-referenced to linked mastoids, filtered (high-pass: 1 Hz, low-pass: 100 Hz, band-stop filter: 48-52 Hz), demeaned and segmented in 2 second epochs for a first visual artifact rejection. All phases of the Memory Arena and the PVT were concatenated, coarse artifacts were removed based on outliers regarding amplitude, kurtosis and variance (implemented in *ft_rejectvisual*) and bad channels were rejected. Based on those data, the unmixing matrix was obtained and bad components were identified. The raw data were then preprocessed again, as the first preprocessing was optimized for ICA. The data were down-sampled to 200 Hz, re-referenced to linked mastoids, filtered (high-pass: 0.3 Hz, low-pass: 40 Hz) and demeaned. The unmixing matrix was applied to the new preprocessed data and bad channels were interpolated.

To derive encoding patterns, the encoding part of the Memory Arena was contrasted against the PVT. Note that the PVT was conducted in temporal proximity to the Memory Arena and requires sustained attention but no memory-related processes. The data recorded during the encoding part of the Memory Arena and the PVT were segmented into 1 second epochs (50% overlap), tapered with a Hanning window and transformed from time to frequency domain using Fast Fourier Transformation. To facilitate reproducibility of results, artifacts were defined based on the 95^th^ percentile uniquely for each frequency bin distribution (across epochs). All 1 second epochs above the 95^th^ percentile were labelled as artifacts and excluded. Note that results did not qualitatively change as a function of the chosen percentile (applying the 90^th^ or 85^th^ percentile as a threshold revealed similar results). Power spectra obtained from encoding and PVT were contrasted (encoding - PVT), yielding absolute power changes during encoding relative to the PVT.

Significant frequency bins were defined based on the group statistics by applying a two-sided cluster-based permutation test with 1000 randomisations ^60^. The topography for each participant was then derived by collapsing power values across the significant frequency bins for each channel separately resulting in a 1 x channel (=58) vector.

#### Event Detection

Sleep spindles and SOs were detected for each participant, based on established detection algorithms ^61,62^. Like wake data, sleep data were down-sampled to 200 Hz, re-referenced to linked mastoids and filtered (high-pass: 0.3 Hz, low-pass: 40 Hz). Bad channels were matched between wake and sleep data, excluded and interpolated. Finally, to identify and mark coarse artifacts, data were visually inspected. Channelwise event detection of both sleep spindles and slow oscillations were conducted on data from non-rapid eye movement (NREM) sleep stages 2 and 3. Events were only included if free of artifacts between 1s before and 1s after the event.

To detect fast sleep spindles, data were band-pass filtered between 12-15 Hz (4^th^ order two-pass Butterworth filter). The envelope of the signal was calculated with a moving average of 200ms. An amplitude criterion (mean + 1.25*SD) was applied to the signal. Sleep spindles were detected when the signal exceeded the amplitude criterion for more than 0.5 but less than 3 seconds (duration criterion).

The maximum of the envelope of each detected spindle was used as the amplitude measure. Duration was the time from beginning to end of each event and density was calculated as the number of detected events / total (artifact free) time spent in NREM sleep stage 2 and 3.

To detect slow oscillations, data were band-pass filtered between 0.3-1.25 Hz (4^th^ order two-pass Butterworth filter). Zero crossings were identified, and three criteria (duration criterion, trough to peak criterion and amplitude criterion) had to be fulfilled. The length criterion was met if one positive to negative crossing was followed by a second positive to negative crossing within a time window of 0.8 to 2 seconds. Based on all sufficiently long events, mean and standard deviation were calculated for trough to peak amplitudes as well as for absolute values of trough amplitudes. All events exceeding both means + 1.25*SDs were considered slow oscillations.

The amplitude of SOs was defined as the most negative trough (downstate). To facilitate comparability between sleep measures, the absolute value of SO amplitudes was used. Thus, the downstate became positively scaled to match all other sleep measures. Duration was defined as the time between the first positive to negative and the following positive to negative crossing. Slow oscillation density was calculated by dividing the number of detected events by the total (artifact free) time spent in NREM sleep stages 2 and 3.

Amplitude, duration and density of detected spindles and SOs were extracted per channel and averaged across events (amplitude, duration) resulting in 6 different 1 x channel (=58) vectors.

#### Coupling of sleep spindles

For each spindle event, the phase of the EEG trace filtered in the SO frequency band was extracted. To this end, the data around each spindle event were filtered from 0.3 – 1.25 Hz. After applying a Hilbert transform, the instantaneous phase angle at the maximum of the envelope of each detected spindle was extracted.

To compare spindles with higher vs. lower coupling, all spindle events per channel were classified based on their phase value. The 50% of spindle events with a phase value closest to 0 degrees were classified as spindles with a higher coupling. The remaining 50% of spindle events were classified as spindles with a lower coupling.

#### Comparison between encoding and sleep pattern

For each participant, the 1×58 vector obtained from encoding was correlated (Spearman’s rho) with every 1×58 vector of the sleep characteristics (amplitude, duration and density for spindles and SOs). Correlations were then Fisher z-transformed for group statistics.

### Statistics

Correlation distributions between encoding and sleep topographies were tested with a 2 (event type: spindles vs. SOs) x 3 (event characteristic: amplitude, duration, density) repeated measures ANOVA. Paired sampled t-tests and one-sample t-tests were used for post-hoc comparisons.

Partial correlations were conducted to test for mediating effects of spindle characteristics (e.g., spindle density) on the correlation between encoding and another spindle characteristic (e.g., spindle amplitude).

To test for an association between sequence retention and the encoding-spindle overlap, the Spearman’s rho correlation was conducted. To rule out that the correlation with behaviour is solely driven by either sleep spindles or by encoding power, we applied a permutation approach and shuffled topographies between participants (1000 permutations). To obtain the observed correlation, we derived, for each participant, (i) behavioural performance, (ii) encoding topography and (iii) sleep spindle topography. By shuffling only one topography (encoding or sleep spindles) between participants while retaining the other participant-specific topography and behavioural performance, two null distributions were generated: First, a distribution under the null hypothesis that the participant-specific encoding topography is irrelevant for the correlation with behaviour (shuffling the encoding topographies). Second, a distribution under the null hypothesis that the participant-specific sleep spindle topography is irrelevant for the correlation with behaviour (shuffling the spindle topographies). The observed correlation was then tested against both null distributions.

## Supplemental Information

**Table S1.** Descriptive sleep data in minutes (mean ± SEM). n = 19. TST = total sleep time.

| N1 | N2 | N3 | REM | TST |
| --- | --- | --- | --- | --- |
| 18.39 | 52.53 | 10.42 | 16.24 | 103.58 |
| $\pm 2.46$ | $\pm 3.66$ | $\pm 2.56$ | $\pm 2.85$ | $\pm 2.28$ |

**Figure S1.**
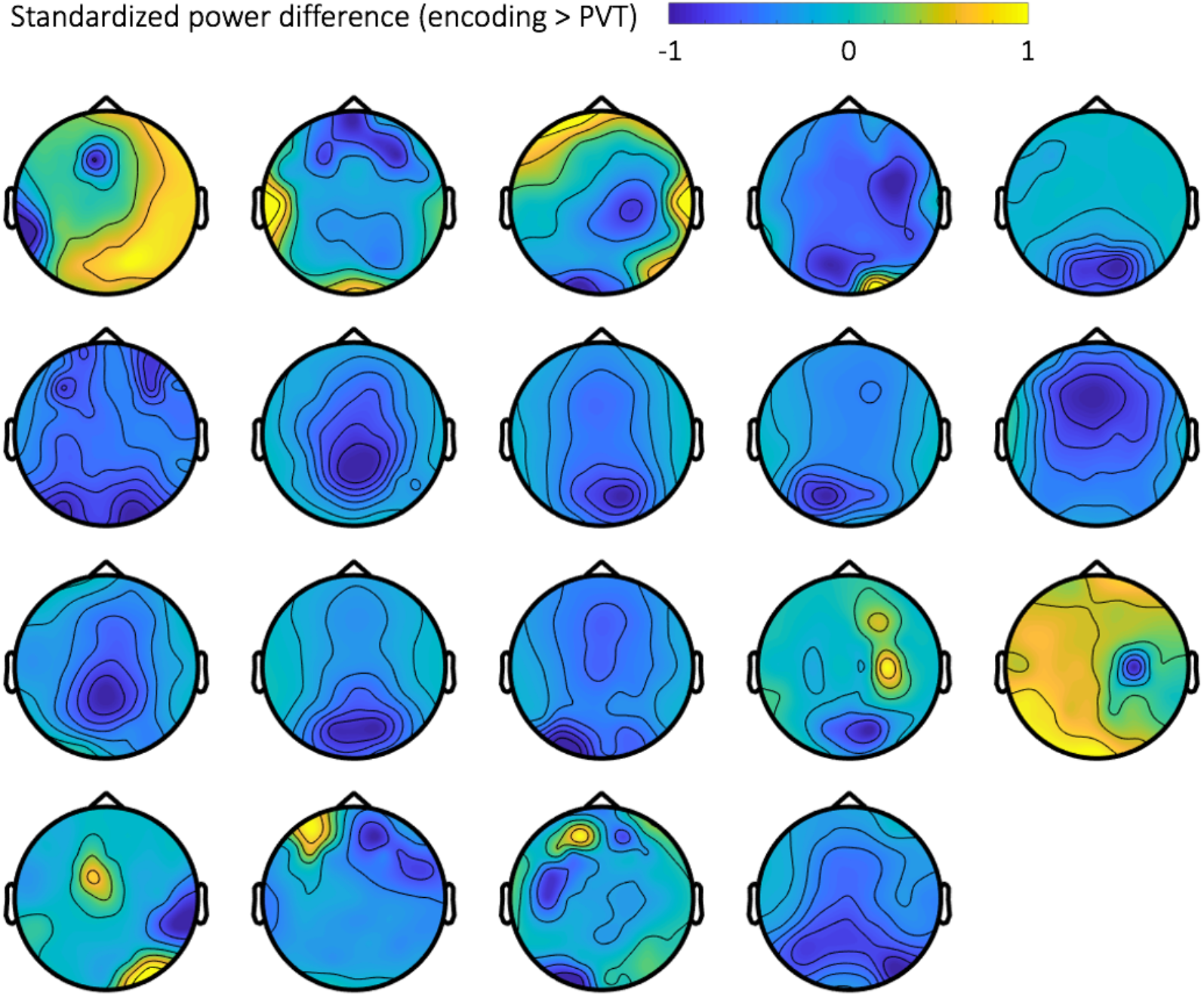
Participant-specific topographies of the 6-20 Hz power changes during encoding relative to the PVT. Power changes were standardized between 1 and −1 for comparability across participants.

**Figure S2.**
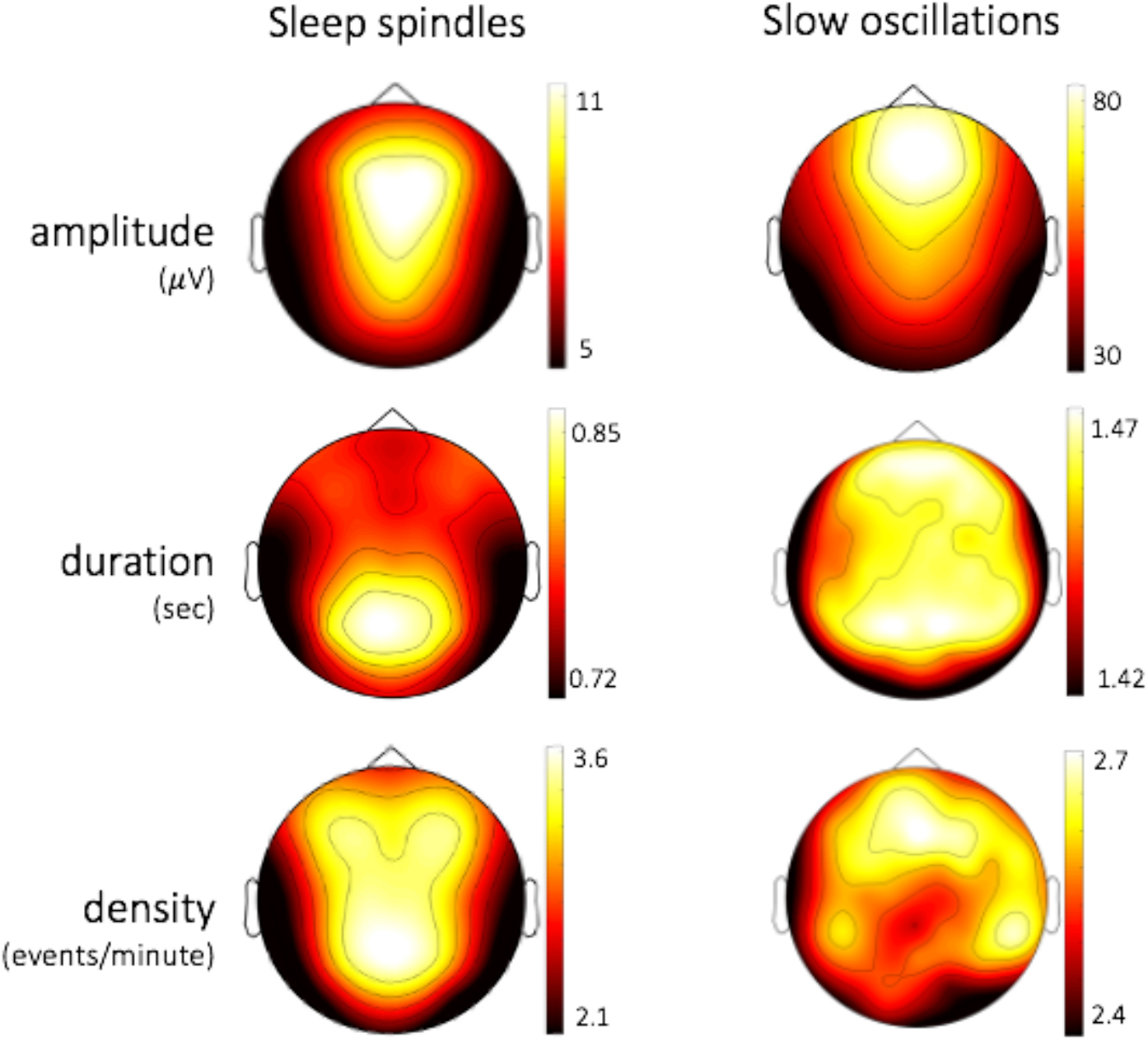
Group-level topographies of amplitude, duration and density of sleep spindles and slow oscillations.

**Figure S3.**
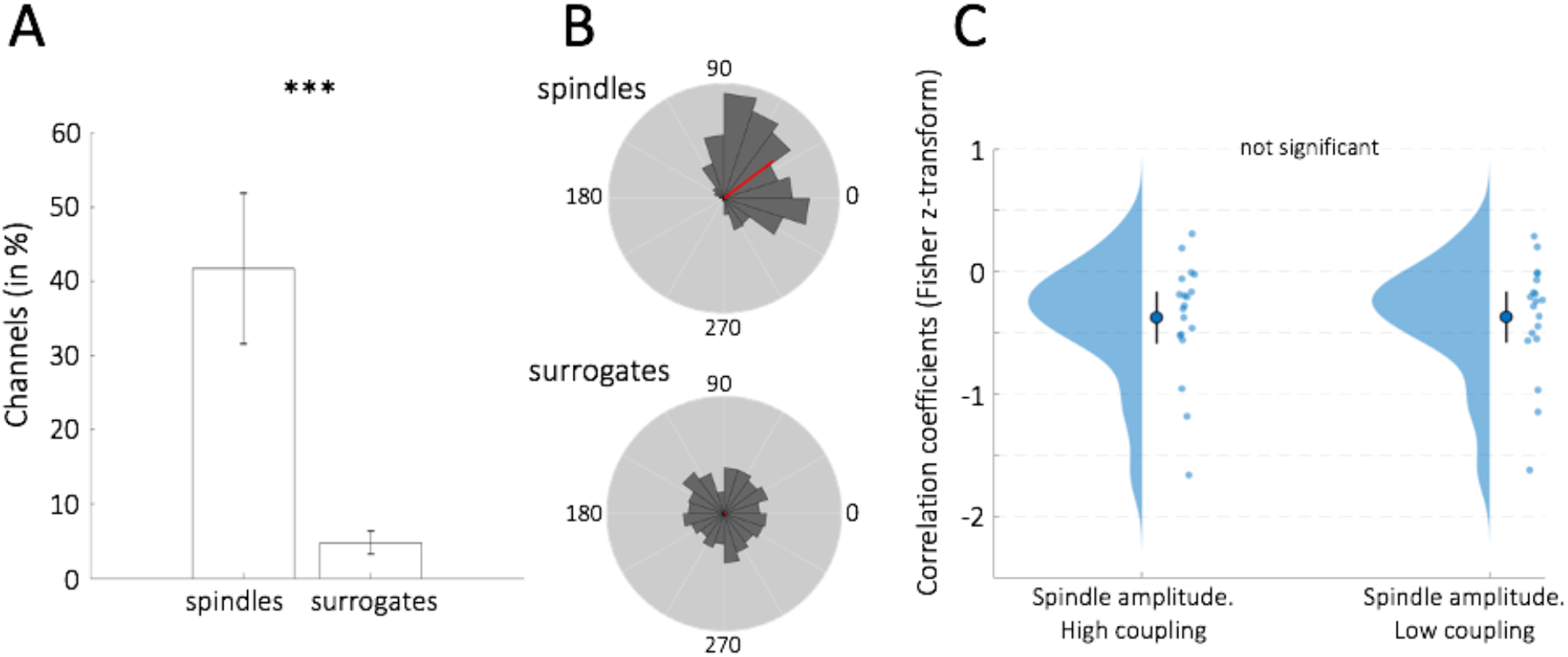
Coupling of spindles and surrogate events to the phase of the signal filtered in the SO frequency range (0.3 −1.25). (A) Mean (+/− 95% confidence intervals) percentage of channels on which sleep spindles and surrogate events are significantly coupled (defined by a significant deviation from a uniform distribution, Rayleigh test: p < .05). Spindles are coupled to SOs on significantly more channels than surrogates. Surrogates were matched control events - for each detected spindle, a spindle-free epoch within 15 seconds before or after the actual spindle event was identified ^48^. The instantaneous phase angle of the SO filtered and Hilbert transformed signal was then extracted at the centre of the spindle-free epoch. (B) The corresponding phase (in degrees) of spindle maxima (top) and surrogate centres (bottom) plotted across all detected events on channels with significant spindle coupling (including all participants, fixed-effects). While spindles significantly cluster at a phase of 37 degrees (Rayleigh test: z = 158.86, p < .001, resultant vector length = 0.59), surrogates do not deviate from a uniform distribution (Rayleigh test: z = 0.94, p = 0.392). (C) No differential encoding-spindle amplitude overlap for spindles with higher (left) vs. lower (right) coupling to the SO up-state.

